# SAFEPPP: a Simple And Fast method to Find and analyze Extreme Points of a metabolic Phenotypic Phase Plane

**DOI:** 10.1101/642363

**Authors:** Mohammad Hossein Moteallehi-Ardakani, Sayed-Amir Marashi

## Abstract

There are many algorithms that help us understand how genome-scale metabolic networks work and what are their capabilities. But unfortunately, the majority of these methods are based on integer linear programming suffer from long run times and high instrumental demand. Optimal solutions in any constraint-based modeling as genome-scale metabolic networks models are on the extreme points of the solution space. We introduce a fast and simple toolbox that reveals extreme characters of metabolic networks in desired situations which can unmask the hidden potentials of metabolic networks. Determining the possibility of coupling between two desired reaction and the capability of synergic substrate consuming are examples of the applications of this method. Fast enumeration of elementary flux modes that exist in extreme points of phase plane of any two reactions is another achievement of this study.

## Introduction

The importance of analysis of the extreme behaviors of a flux space has been discussed extensively in the literature [16]. Edwards et al. [3] have shown that under optimal growth condition, the metabolic behavior of *Escherichia coli* lies on a special extreme of flux space called “line of optimality (LO)”. Interestingly, when *E. coli* is grown on a new unconventional carbon source, it takes several generations to adapt to the new conditions, but it finally finds the optimal strategy. That is, to adapt its behavior on the marginal location of the LO [7].

Elementary flux modes (EFMs) are minimal set of reactions which can support a feasible steady state condition [13]. Enumeration of Elementary flux modes (EFMs) has been the aim of several recent studies [18,6,2]. EFMs are non-decomposable strategies of metabolism that lie on the extreme points of the flux space and help us understand the extreme behaviors of the metabolic model. An extreme point of a metabolic flux space is more informative than its surrounding points because non-extreme points can be written as a convex combination of extreme points (see Appendix A). In this paper, two kinds of extreme point will be considered.

- First, extreme points in the phase plane of two reactions which show existence of at least one elementary mode that contains the reactions (Figure 1). Such extreme points demonstrate the feasibility of coupling between reactions [8]. Coupling attracts great attention due to the importance of biomass coupled product synthesis [19]. If an EFM exists such that it shows a coupled production of both biomass and a desired product, then there exists a knockout strategy to enforce the organism to produce the desired product as the obligate byproduct of growth [8].

**Figure 1.**
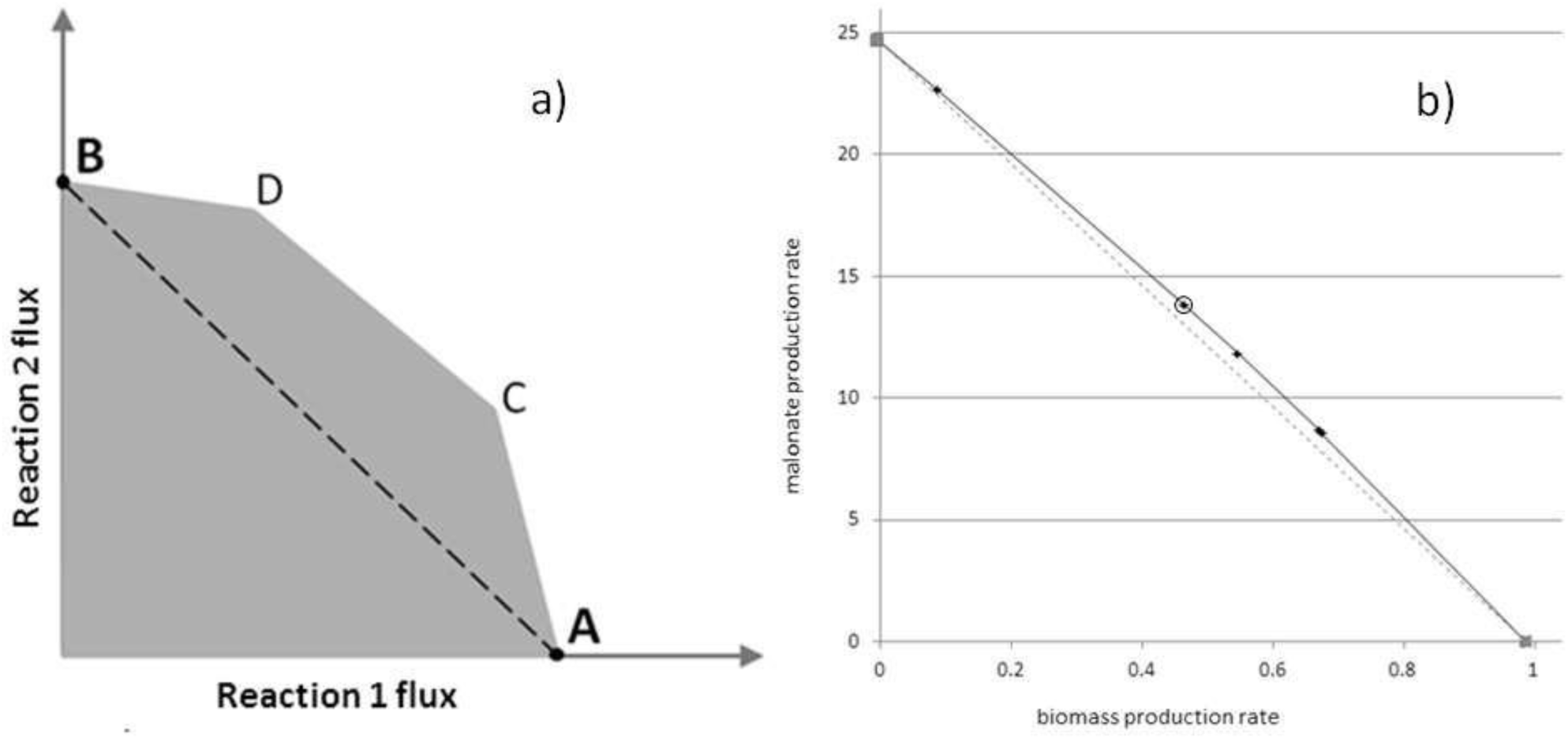
(a): Phase plane in our case is the projection of flux space on the plane of two specific reactions. *E. coli* metabolic model when consuming 1 mole of substrate has a maximum flux of **A** and **B** for reactions 1 and 2, respectively. All flux vectors whose projection lies on the dashed line can be obtained as a convex combination of A and B. The phase plane beyond the dashed line shows that the network can simultaneously maximize linear combination of reactions 1 and 2 fluxes more efficiently than using the sum of extreme strategies A and B. All of the borders of phase plane can be written as convex combination of extreme points (A, B, C, D). In other words, the extreme points are non-decomposable maximal possible strategies of the network. Each of these points is the projection of at least one EFM on phase plane. (b): an example of a real phase plane. Reactions 1 and 2 are biomass and malonate production rates, respectively. There are seven extreme points in this phase plane, five of which can produce biomass and malonate simultaneously. The circled extreme point includes an eight reactions minimal knockout strategy (MKS) that couples malonate production with growth. Minimal knockout strategies of the other extreme points include more reactions.
- Second, extreme points of the plot of biomass production against different substrates. Such an extreme point shows a growth condition in which by synergistically consuming two different substrates, the organism can produce more biomass than the sum of biomass produced by each substrate alone.

## Software description

Determining the phase plane, elementary modes and knock out strategies have many applications as mentioned previously. Here we introduce a software that gets model and desired reactions and gives these three above mentioned as output. All functions are compatible with the COBRA toolbox environment in Matlab. Here we first shortly explain the previously-introduced functions which are used in our code.

### FBA

FBA can maximize any linear combination of fluxes under constraints of the problem. The mathematical representation of FBA is [14]:

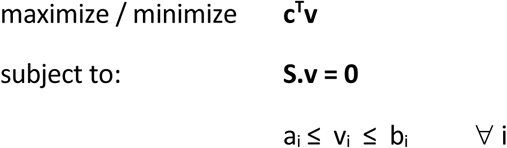

### Flux coupling analysis

Similar to Tabe-Bordbar, Marashi [18] we use F2C2 tool [9] for a more rational reaction elimination. F2C2 gets the model as the input and returns two variables, “blocked” and “fctable”. “blocked” is the set of blocked reactions that cannot have any non-zero flux in any steady state condition, while “fctable” is an N×N matrix (N: number of unblocked reactions) in which determines the coupling relation of each reaction pair. The value of fctable(i,j) is 0, 1 and 2 when reactions i and j are uncoupled, fully coupled and partially coupled respectively; and is 3 and 4 when reaction i is directionally coupled to j and reaction j is directionally coupled to i, respectively.

A number of functions are developed in this study:

### ExPo

This function finds the extreme points of phase plane in a recursive manner (Figure2). The extreme points were found so fast and efficiently.

**Figure 2:**
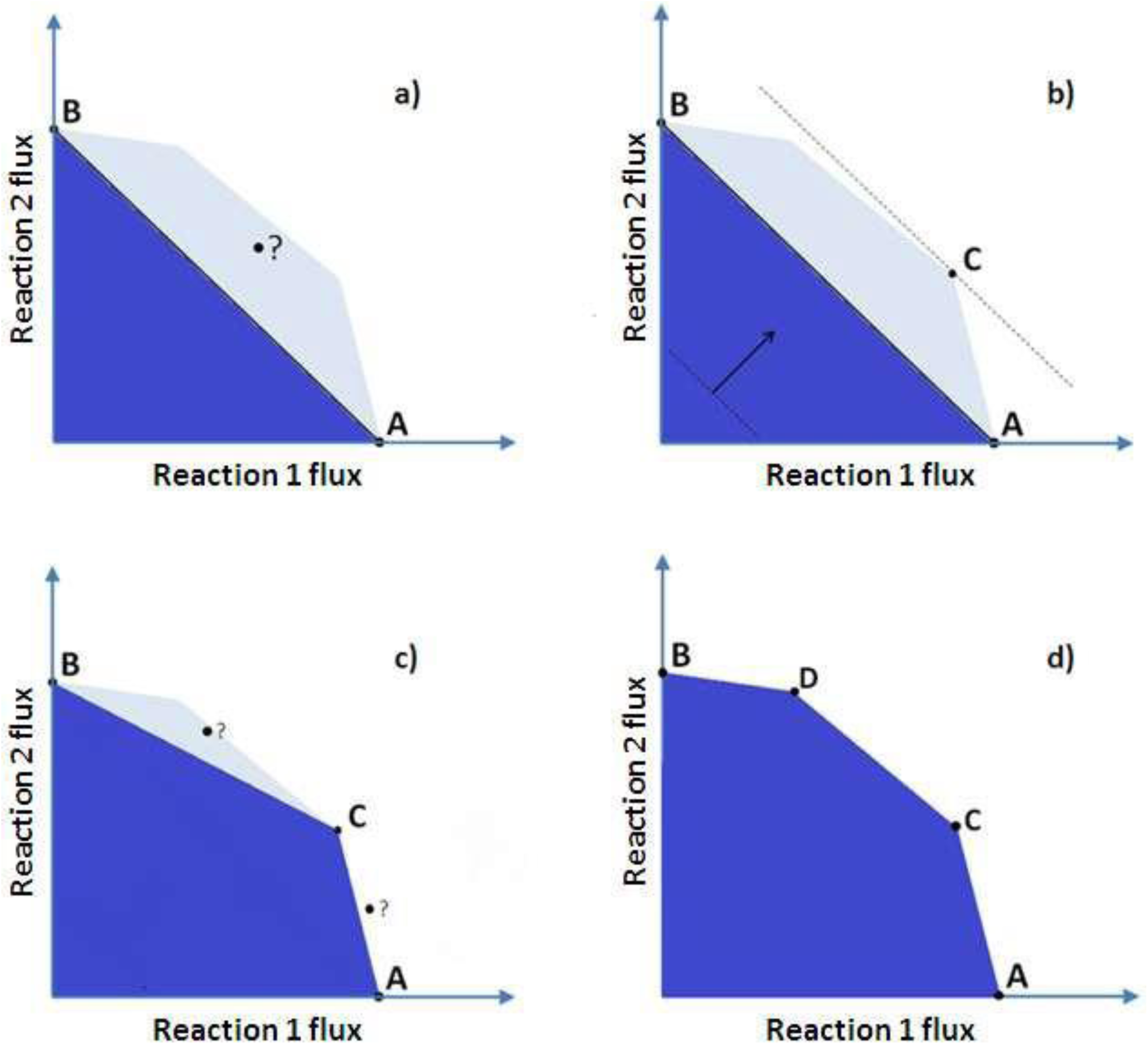
(a) at the first step, **A** and **B** were found by FBA when reaction 1 and reaction 2 were set as objective functions respectively. The discovered portion of phase plane is dark blue. At this step, the algorithm tries to find an extreme point beyond the A-B line; (b) Here the algorithm calls a recursive function, RExPo(A,B), which determines the objective function of FBA such that it ensures us that any extreme point beyond the A-B line is found (if any). At this step the algorithm finds **C**; (c) Now there are two new problems like the previous problem. The algorithm calls RExPo(A, C) and RExPo(C, B) to answer the questions. (d) There is an extreme point between B and C, **D**. the algorithm continue to search extreme points between founded points, but there is no more extreme points. So the function returns these four extreme points.

### FindEM

Following the approach of Tabe-Bordbar, Marashi [18], FindEM function aims to determine the elementary flux modes that lie on one specific extreme point by a “random elimination” procedure. The function randomly removes K active reactions of the extreme point flux vector at each step and checks the feasibility of the extreme point in the reduced metabolic model. If the extreme point becomes infeasible, the removed group of reactions will be restored to the model; otherwise, the deletion is accepted. Random elimination continued till achieving a non-reducible subset of reactions that allow steady-state condition at the extreme point, which is an EFM. The value of K is updated in each step in such a way that maximizes the expected value of number of accepted deleted reactions (see Appendix B).

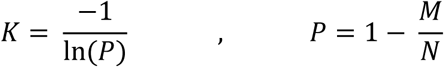

Where M is the mean number of reactions in each EFM and N is the number of undeleted reactions in the removable reaction list of the algorithm at each step.

### RARR (Reactions that are Absolutely Required to be Removed)

This function determines the set of reactions that are required to be removed in any knockout strategy for coupling. Another version of the function, Drarr, determines the set of reaction pairs that at list one should be removed.

### MKSF (Minimal Knockout Strategy Finder)

EFMs that make the flux vector of specific extreme points have non-zero fluxes for both reactions. Non-reducibility of EFMs ensures us that these two reactions can be coupled. Coupling is especially interesting when one of the reactions is the production of desired product, and the other is a cellular objective function. An important example is ‘biomass coupled product synthesis’ [8]. Existence of extreme points at the phase plane of biomass and a desired product means that there is a knockout strategy which causes the cell to grow only at the expense of desired product synthesis. A trivial strategy to achieve coupling is removing all inactive reactions of the extreme points from the metabolic model. It always works in theory, but it is not applicable in reality; because extensive manipulation, in practice, harms the viability of the cell.

Finding minimal knockout strategies is the aim of MKSF function. In this case, ‘random elimination procedure’ eliminates K reactions from inactive reaction list of the extreme point (activate them) and check for coupling by FBA. If by activating a group of reactions, growth and the production of the desired product becomes uncoupled, the group will be returned to the inactive reaction list. K is determined the same as before.

### RRE (rational reaction elimination)

The main strategy of this work is random inactivation of active reactions of an extreme point till achieving an EFM and also random activation of inactive reactions of an extreme point till achieving an MKS for coupling. To reduce the problem size before using the random elimination procedure, we use rational reaction elimination (RRE) which eliminates the following reactions:

- Blocked reactions: Blocked reactions are not a member of any EFM, and also their inactivation is unnecessary for any knockout strategy.
- Fully coupled reactions: in any group of fully coupled reactions, activation or inactivation of each member enforce the entire group to be activated or inactivated respectively. Therefore, to reduce the problem size, the algorithm eliminates all fully coupled reactions except one member of each group.
- Essential reactions: To find a full set of EFMs on an extreme point and to find all possible knockout strategies of an extreme point, the algorithm eliminates essential reactions. Essential reactions are reactions that are present in all possible answers. In the case of finding EFMs essential reactions are reactions that are present in all EFMs of extreme point and so are necessary for the feasibility of extreme point. In the case of finding MKSs, essential reactions are reactions whose sole activation can uncouple our two desired reactions.

RRE also reduces the size of the problem by setting:

- Elimination priorities: Three above mentioned reactions are eliminated before the random elimination procedure but “setting elimination priorities” guides the random elimination procedure. The algorithm puts the remaining reactions in priority groups. The highest priority group is a group of reactions that enforces more reaction elimination due to directionally coupled relations between reactions (see Figure1 in reference [18]). The random elimination procedure starts with high priority groups.

### SYN

This function determines the synergy patterns of two substrates on maximizing the flux of an objective reaction, usually biomass production (Figure 3). The extreme points of the plot are related to at list one EFM. These extreme points show us the possibility of designing strains which co-utilize two substrates [4] because these are non-decomposable. In other words, it means that there exists a strategy to make the two substrate uptake reactions coupled.

**Figure 3:**
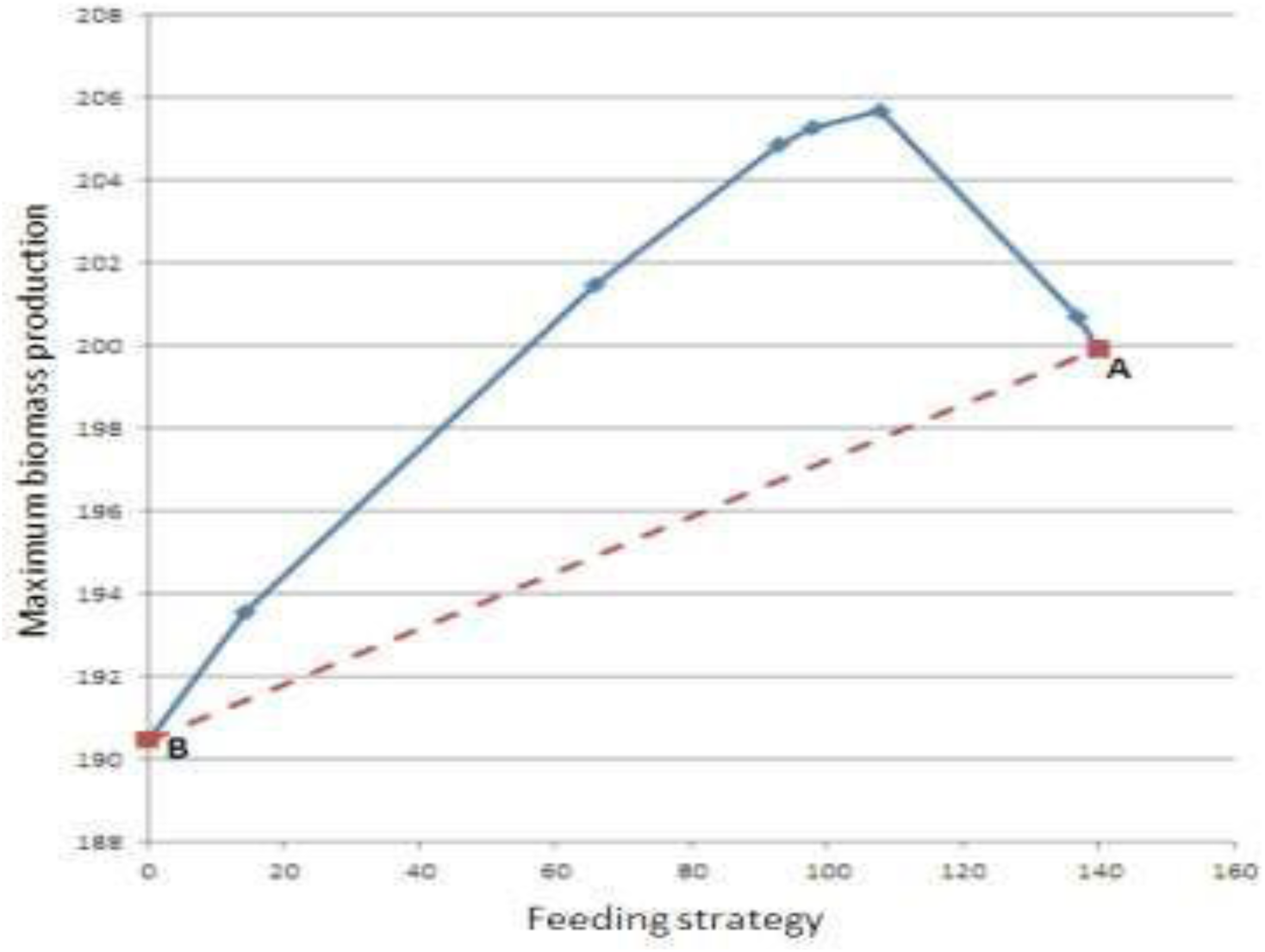
The synergistic effects of consuming both glucose and prolyl-Glycine on the growth of *E. coli* metabolic model are shown. The algorithm first determines the biomass production attainable by consuming glucose or prolyl-Glycine lonely, A and B respectively. If there not exists any synergy, we expect the maximum biomass to be on convex combination of points A and B (dashed line), but when we give combinational feeding to the model, we see more biomass production. In this case, 140 feeding conditions (*N* = 140) tested by the algorithm, two pure and 138 combinational feedings. *N* is determined by the user and specifies the precision of the algorithm (small values of *N* may lead the algorithm to find two adjacent/near extreme points as one). The location of extreme points doesn’t depend on *N*.

### Implementation

All optimization problems were solved by ‘’glpk” [12]. As for the algorithm, all the data and operations other than solving optimization models were processed in the COBRA toolbox environment [17]. All computations were performed in a PC with a 2.67GHz CPU and 24 GB of RAM.

## Applications

Fast determination of phase plane of any two reactions, is one the most important applications of the software which is necessary for phase plane analysis [11,10,5]. The phase plane of biomass and ethanol in *E. coli* under anaerobic condition (in which O2 uptake is blocked) is shown in Figure4.

**Figure 4:**
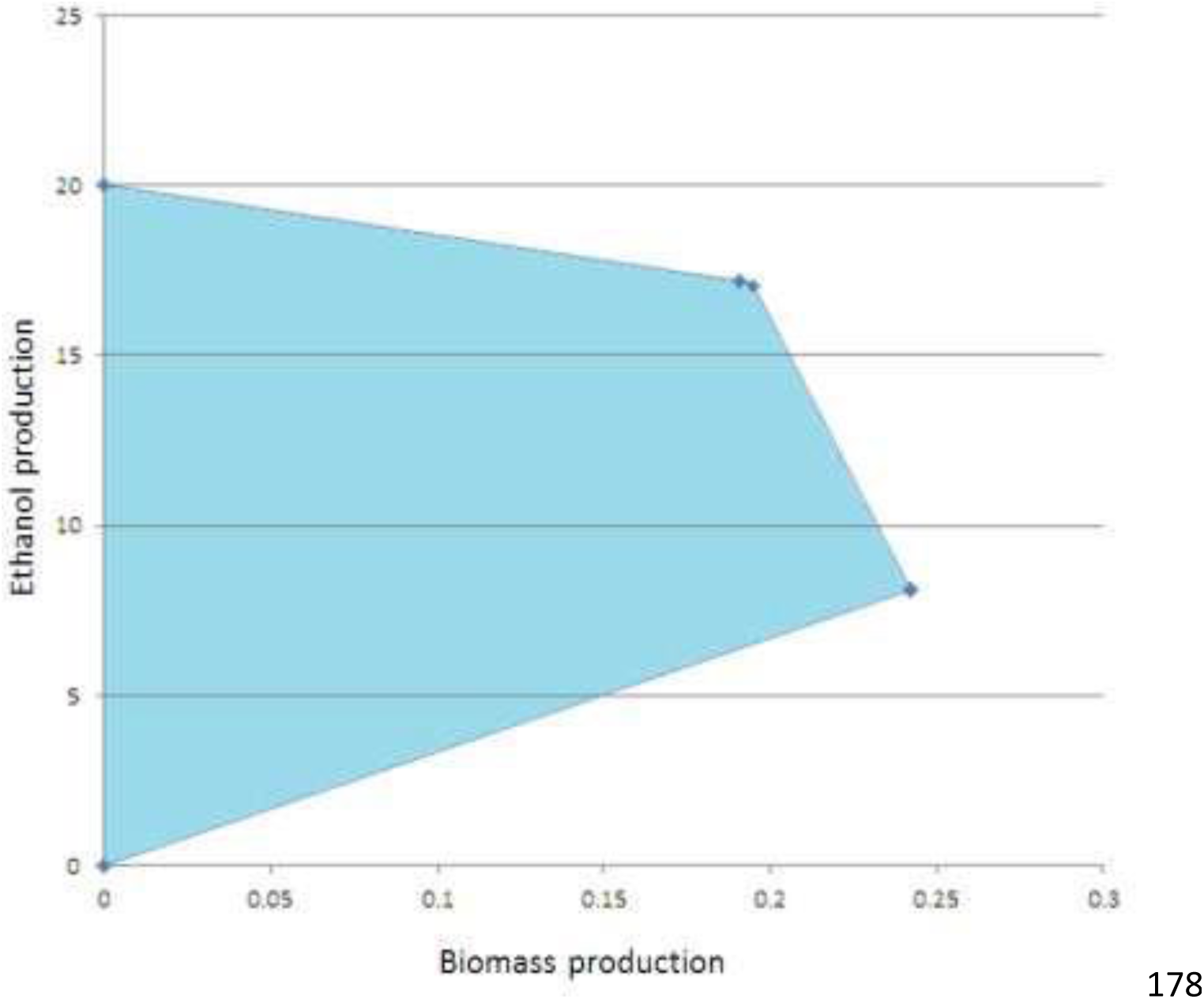
phase plane of biomass an ethanol production (production envelope) under anaerobic condition. The phase plane shows ethanol production in maximum biomass production, and it is consistent with our knowledge of anaerobic growth of *E. coli*.

The phase plane of biomass and all other active reactions (1389 reactions with maximum flux of more than 0.1) was determined in an average time of 0.83 second (due to complexity of the biomass reaction when the phase plane is plotted as biomass production versus production of a certain metabolite, computation is generally slow. However, when the phase plane of production of two metabolites is plotted, computation is much faster. 601 reactions don’t have any nontrivial extreme points. And there is one reaction in the model with 12 nontrivial extreme points (Figure5).

**Figure 5:**
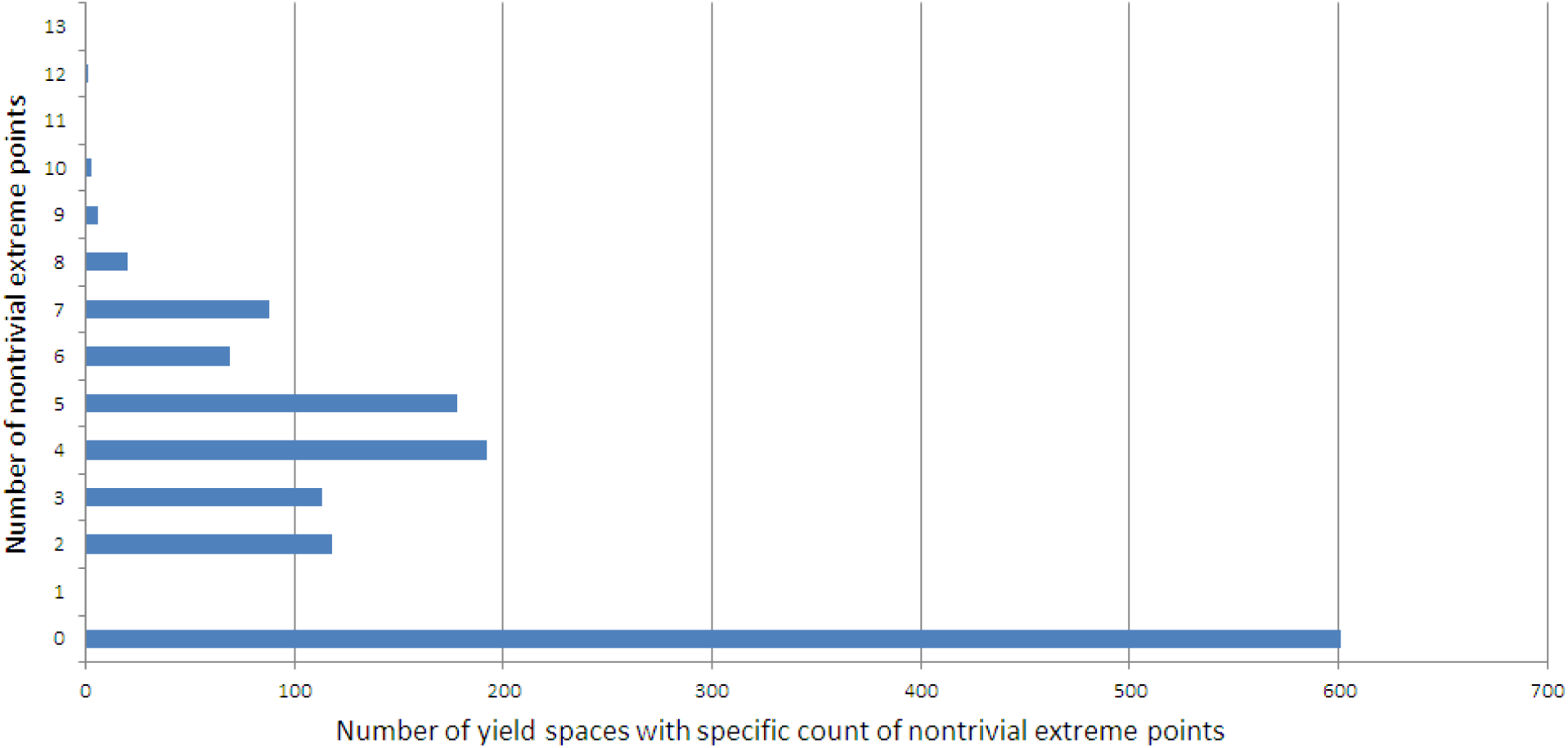
distribution of active reactions against their number of non-trivial extreme points in their biomass phase plane. The maximum number of nontrivial extreme point is 12.

In Figure1 (b), we mention to an example of minimal knockout strategies (MKS) found by our software which couples malonate and biomass production. The knockout strategy has eight reactions and, to our knowledge, it is the first reported knockout strategy to couple biomass and malonate production. Because of the increasing complexity of the problem, finding knockout strategies of five reactions and more is not applicable by algorithms like OptKnock[1] and OptGene[15]. Due to randomness of the algorithm, new MKSs founded harder and harder in time (Figure 6).

**Figure 6:**
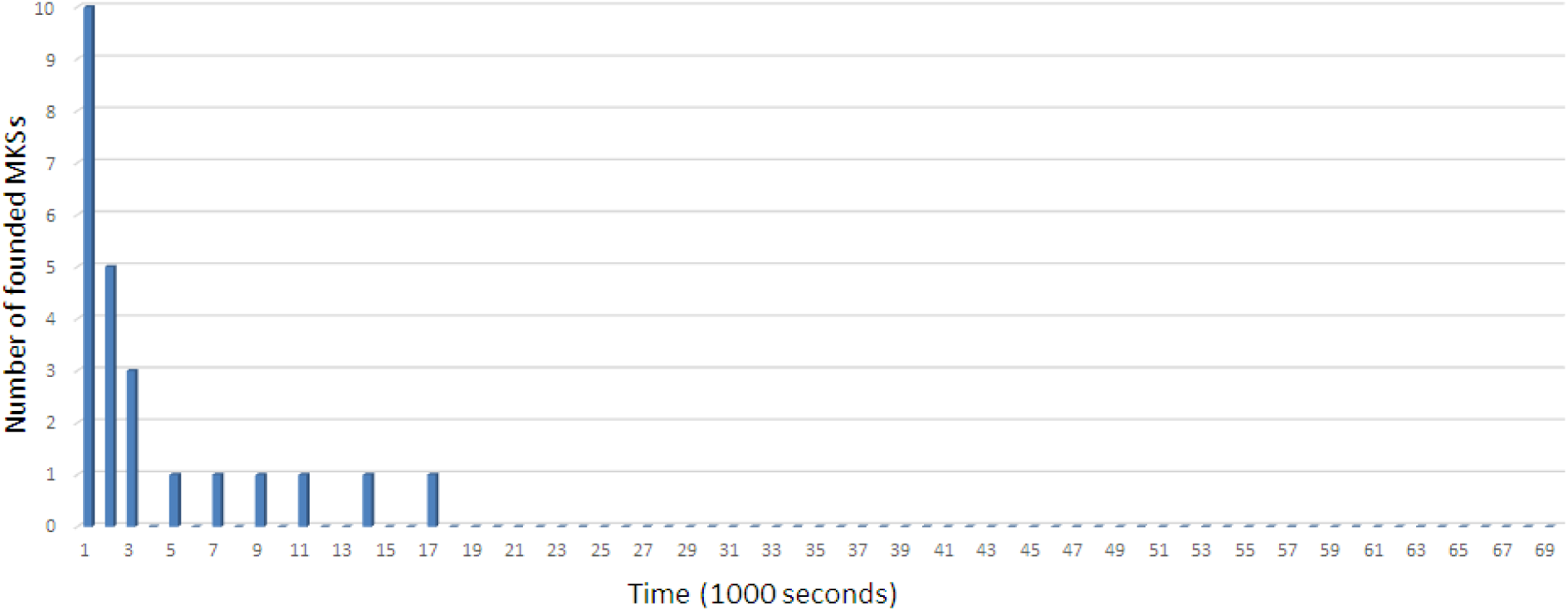
time distribution of 24 MKSs founded in 69000 seconds. The smallest MKS has eight reactions. Each MKS was founded in less than one minute.

## Acknowledgment

We are thankful to Dr. S. Asad (University of Tehran) because of her contribution to developing the idea of this work and also F. Marefat (Amirkabir University) for her help on scripting the code.

### Appendix A

#### Proposition

Existence of an extreme point of phase plane means the existence of at least one elementary mode which it is projected on the extreme point.

*Proof*: Suppose that the opposite is correct. Assume that the flux vector, whose projection is on the mentioned extreme point, is composed of more than one elementary mode whose projections are not on the same extreme point. It means that the flux vector can be written as a convex combination of some elementary modes. All of the elementary modes have their own projected point on the phase plane. An extreme point of a convex shape cannot be written as a convex combination of the other points of the shape. So the assumption cannot be correct.

### Appendix B

Consider *M* as the mean number of reactions in each EFM and *N* as the number of undeleted reactions in the removable reaction list of the algorithm at each step. So 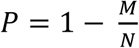 is the probability of accepting one reaction deletion. The probability of accepting K reaction deletion can be estimated by 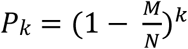. So the expected value of ‘number of accepted deleted reactions’ after a K deletion is *E*[*K*] = *K* × *P*_*k*_ The value of K that maximize the E[K] is 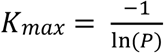

## References

1. Burgard AP, Pharkya P, Maranas CD (2003) Optknock: a bilevel programming framework for identifying gene knockout strategies for microbial strain optimization. Biotechnology and Bioengineering 84:647–657

2. Chan SHJ, Ji P (2011) Decomposing flux distributions into elementary flux modes in genome-scale metabolic networks. Bioinformatics 27:2256–2262

3. Edwards JS, Ibarra RU, Palsson BO (2001) In silico predictions of *Escherichia coli* metabolic capabilities are consistent with experimental data. Nature Biotechnology 19:125–130

4. Gawand P, Hyland P, Ekins A, Martin VJ, Mahadevan R (2013) Novel approach to engineer strains for simultaneous sugar utilization. Metabolic Engineering 20:63–72

5. Horvat P, Koller M, Braunegg G (2015) Recent advances in elementary flux modes and yield space analysis as useful tools in metabolic network studies. World Journal of Microbiology and Biotechnology 31:1315–1328

6. Hunt KA, Folsom JP, Taffs RL, Carlson RP (2014) Complete enumeration of elementary flux modes through scalable, demand-based subnetwork definition. Bioinformatics:btu021

7. Ibarra RU, Edwards JS, Palsson BO (2002) *Escherichia coli* K-12 undergoes adaptive evolution to achieve in silico predicted optimal growth. Nature 420:186–189

8. Klamt S, Mahadevan R (2015) On the feasibility of growth-coupled product synthesis in microbial strains. Metabolic Engineering 30:166–178

9. Larhlimi A, David L, Selbig J, Bockmayr A (2012) F2C2: a fast tool for the computation of flux coupling in genome-scale metabolic networks. BMC Bioinformatics 13:57

10. Lopar M, Špoljarid IV, Atlid A, Koller M, Braunegg G, Horvat P (2013) Five-step continuous production of PHB analyzed by elementary flux, modes, yield space analysis and high structured metabolic model. Biochemical engineering journal 79:57–70

11. Lopar M, Špoljarid IV, Cepanec N, Koller M, Braunegg G, Horvat P (2014) Study of metabolic network of Cupriavidus necator DSM 545 growing on glycerol by applying elementary flux modes and yield space analysis. Journal of industrial microbiology & biotechnology 41:913–930

12. Makhorin A (2008) GLPK (GNU linear programming kit).

13. Marashi S-A, David L, Bockmayr A (2012) Analysis of metabolic subnetworks by flux cone projection. Algorithms for Molecular Biology 7:17

14. Orth JD, Thiele I, Palsson BØ (2010) What is flux balance analysis? Nature Biotechnology 28:245–248

15. Patil KR, Rocha I, Förster J, Nielsen J (2005) Evolutionary programming as a platform for in silico metabolic engineering. BMC Bioinformatics 6:1

16. Raman K, Chandra N (2009) Flux balance analysis of biological systems: applications and challenges. Briefings in Bioinformatics 10:435–449

17. Schellenberger J, Que R, Fleming RM, Thiele I, Orth JD, Feist AM, Zielinski DC, Bordbar A, Lewis NE, Rahmanian S (2011) Quantitative prediction of cellular metabolism with constraint-based models: the COBRA Toolbox v2. 0. Nature protocols 6:1290–1307

18. Tabe-Bordbar S, Marashi S-A (2013) Finding elementary flux modes in metabolic networks based on flux balance analysis and flux coupling analysis: application to the analysis of *Escherichia coli* metabolism. Biotechnology letters 35:2039–2044

19. Toya Y, Shiraki T, Shimizu H (2015) Ssdesign: Computational metabolic pathway design based on flux variability using elementary flux modes. Biotechnology and Bioengineering 112:759–768

